# *In silico*-based screening for natural product’s structural analogs as new drugs candidate against leishmaniasis

**DOI:** 10.1101/2022.07.22.501189

**Authors:** Haruna Luz Barazorda-Ccahuana, Luis Daniel Goyzueta-Mamani, Mayron Antonio Candia-Puma, Camila Simões de Freitas, Grasiele de Sousa Vieria Tavares, Daniela Pagliara Lage, Eduardo Antonio Ferraz Coelho, Miguel Angel Chávez-Fumagalli

## Abstract

Leishmaniasis is a disease with high mortality rates and approximately 1.5 million new cases each year. Despite the new approaches and advances to fight the disease, there are no effective therapies. Hence, this study aims to *in silico* screen for natural product’s structural analogs as new drugs candidate against leishmaniasis. We applied *in silico* analysis, such as virtual screening, molecular docking, molecular dynamics simulation, and Molecular Mechanics-Generalized Born Surface Area MM/GBSA estimation aiming to select structural analogs from natural products that have shown antileishmanial activity against arginase (ARG) enzyme and that could bind selectively against Leishmania ARG. The compounds 2H-1-Benzopyran, 3,4-dihydro-2-(2-methylphenyl)-(9CI), Echioidinin, and Malvidin showed good results against ARG targets from three parasite species and negative results for potential toxicities. The Malvidin ligand generated interactions in the active center at pH 2.0 conditions and hydrogen bonds enhancing receptor-ligand coupling. This work identified Malvidin as a potential drug candidate to treat leishmaniasis.

## Introduction

Leishmaniasis is an ancient disease that has been described in archaic ceramics, statues, and writings; and, in molecular findings from mummified human bodies and archaeological material [1]. The disease causes high morbidity and mortality worldwide, where about one billion people are at risk of infection across 98 countries worldwide, with over 1.5 million new cases and 20,000–40,000 deaths reported each year [2,3]. The increase in leishmaniasis incidence and prevalence is mainly attributed to several risk factors that are man-propelled [4], whereas, in many regions, the transmission pattern shows expansion with new territories affected by the disease [5,6]. Also, Leishmaniasis has gained greater importance in HIV-infected patients, as an opportunistic infection in areas where both pathogens are endemic [7].

Leishmaniasis is caused by protozoan parasites of the genus *Leishmania* (Kinetoplastida: Trypanosomatidae), which has a digenetic life cycle alternating between a mammalian host and insect vectors [8]; while about 21 parasite species can infect mammals, and many of them cause human disease [9]. The clinical manifestations depend on both the parasite species and the hosts’ immune response [10]; varying from a chronic, slow-to-heal disease known as Tegumentary Leishmaniasis (TL), to a potentially fatal form of the disease in which parasites disseminate to internal organs, such as the liver, spleen, and bone marrow, namely, Visceral Leishmaniasis (VL) [11]. Despite significant progress, the development of a human vaccine remains hampered by significant gaps in the development pipeline [12]; and the treatment against disease has used drugs that cause side effects in the patients, such as myalgia, arthralgia, anorexia, fever, and urticarial, besides toxicity in the liver, kidneys, and spleen [13]. Therefore, the need for cost-effective treatment that completely cures the disease with minimal side effects, low relapse rate, high efficacy, and less toxicity remains to be clarified [14].

The number of natural product-derived drugs present in the total amount of drug launchings in the market over four decades represents a significant source of new pharmacological entities [15]; while a series of secondary plant-purified products has already been described with leishmanicidal potential [16–19]. Likewise, bioinformatics and computational approaches became crucial for rapid *in silico* screening of potential metabolite databases from natural sources that can be repurposed against diseases for faster, safer, and cheaper drug development [20,21]; the alternative strategy of target-based drug discovery is used extensively by the pharmaceutical industry and has been applied to Leishmaniasis [22,23]. However, *in silico* methods to identify new potential drugs to be applied against Leishmaniasis, present limitations, such as the dependency on the quality, accuracy, and completeness of the information present in databases [24].

The arginase (ARG) enzyme has recently received considerable attention for its role in the disease since recent studies have highlighted it as a potential therapeutic target [25]. ARG is the first enzyme of the polyamine pathway in *Leishmania* and catalyzes the conversion of L-arginine to L-ornithine and urea, down-regulating the polyamine pathway, affecting the parasite growth and infectivity [26]. The polyamine pathway inhibition results in the lack of defense against reactive oxygen species generation (ROS) produced by macrophages, degrading the genetic material of the parasite and causing its death by apoptosis [27]. Hence, natural products (NPs) such as glycoside compounds, polysaccharides, and phenolic compounds have shown an effect against arginase activity [28,29]; however, several of these compounds have also shown high affinity to human ARG [30]. Taking this into consideration, in the present work, we used a virtual drug screening based on docking prediction, molecular dynamics simulations, and ligand-binding affinities, aiming to select structural analogs to NPs that have shown antileishmanial activity; as well as active against ARG and that could bind selectively against *Leishmania* ARG, with the purpose to prospect a future therapeutic candidate to be applied against leishmaniasis.

## Results

### Data collection and virtual screening

In this work, a search was performed in the NuBBE database for NPs that has been described with antileishmanial and anti-ARG activity. The search in the database resulted in 33 NPs described with antileishmanial activity, whereas six of them had also been described as inhibitors of ARG activity. Startlingly, all the NPs selected were described in the same article, in which the compounds were isolated from *Byrsonima coccolobifolia* species and tested for *in vitro* anti-ARG activity [31]. Since no antileishmanial activity is reported in the article, a cross-reference search for each compound was performed in the Pubmed database to validate the properties. Thereafter, the SMILEs from Quercetin (NuBBE_122), Isoquercetin (NuBBE_123), Quercitrin (NuBBE_161), (+)-Syringaresinol (NuBBE_214), Catechin (NuBBE_287) and (-)-epicatechin (NuBBE_866) were obtained from PubChem and submitted to physicochemical properties analysis related to Absorption, Distribution, Metabolism, and Excretion (ADME) profile; where Lipinski’s rule of five (MW, iLOGP, HBAs and HBDs) [32], the Quantitative Estimate of Druglikeness (TPSA, RB, NHA and the number of alerts for undesirable substructures) [33] and the Synthetic Accessibility [34], of the NPs are shown in Table 1.

**Table 1.** Natural compounds description selected in the NuBBE database

| NuBBE ID | PubChemID | Name | MW | iLOGP | TPSA | HBA | PAINS | Brenk | SynAcce |
| --- | --- | --- | --- | --- | --- | --- | --- | --- | --- |
| NuBBE_122 | 5280343 | Quercetin | 302.24 | 1.63 | 131.36 | 7 | 1 | 1 | 3.23 |
| NuBBE_123 | 5378597 | Isoquercetin | 464.38 | 2.11 | 210.51 | 12 | 1 | 1 | 5.32 |
| NuBBE_161 | 5353915 | Quercitrin | 448.38 | 1.27 | 190.28 | 11 | 1 | 1 | 5.28 |
| NuBBE_214 | 100067 | (+)-Syringaresinol | 418.44 | 3.52 | 95.84 | 8 | 0 | 0 | 4.36 |
| NuBBE_287 | 1203 | Catechin | 290.27 | 1.47 | 110.38 | 6 | 1 | 1 | 3.50 |
| NuBBE_866 | 72276 | (-)-epicatechin | 290.27 | 1.47 | 110.38 | 6 | 1 | 1 | 3.50 |

MW: Molecular weight; iLOGP: octanol/water partition coefficient; TPSA: Topological Polar Surface Area; HBA: Number of H-Bond acceptors; HBD: Number of H-bond Donors; RB: Rotatable bonds; NHA: Number of heavy atoms; SynAcce: Synthetic accessibility

To find structural analogs to the six NPs selected, a search at the SwissSimilarity server employing the commercial Zinc-drug like compound library was performed, resulting in 400 analogs for each NP; however, the search comprised a high degree of redundancy between the analogs and a step in which duplicated compounds were excluded was executed, resulting in a total of 1499 unique compounds selected for virtual screening (Figure 1). The virtual screening results against *Leishmania infantum* and Human ARG are plotted in Figure 1A, where a positive linear relationship between the binding affinities of the compounds toward both targets is shown [Pearson r: 0.931; r^2^: 0.868]. Later, aiming to select compounds that showed higher affinity toward *L. infantum* ARG, the maximum cut-off values were selected considering the results of the six original NPs, resulting in 25 compounds selected (Figure 1A). Since *in vitro* evidence of inter-species differences in the susceptibility of parasites to antileishmanial drugs have been reported [35], putative drug candidates must be active against several species of the parasite [36]; in this way, to compare the bindings affinities of the 25 selected compounds, results from *L. infatum, L. mexicana* and *L. braziliensis* ARG were normalized against human homologs and plotted in a heatmap, while each compound results showed differences in their normalized affinities profile (Figure 1B). Also, to select potential nontoxic candidates, the tumorigenic, mutagenic, reproductive effect, and irritant action were assessed for the 25 compounds (Figure 1C). Thus, the compounds 2H-1-Benzopyran, 3,4-dihydro-2-(2-methylphenyl)-(9CI) (ZINC39120134) (Figure 1D), Echioidinin (ZINC14807307) (Figure 1E), and Malvidin (ZINC897714) (Figure 1F) were selected for further analysis, since they showed higher normalized affinities against the three parasite species targets and negative results for potential toxicities.

**Figure 1.**
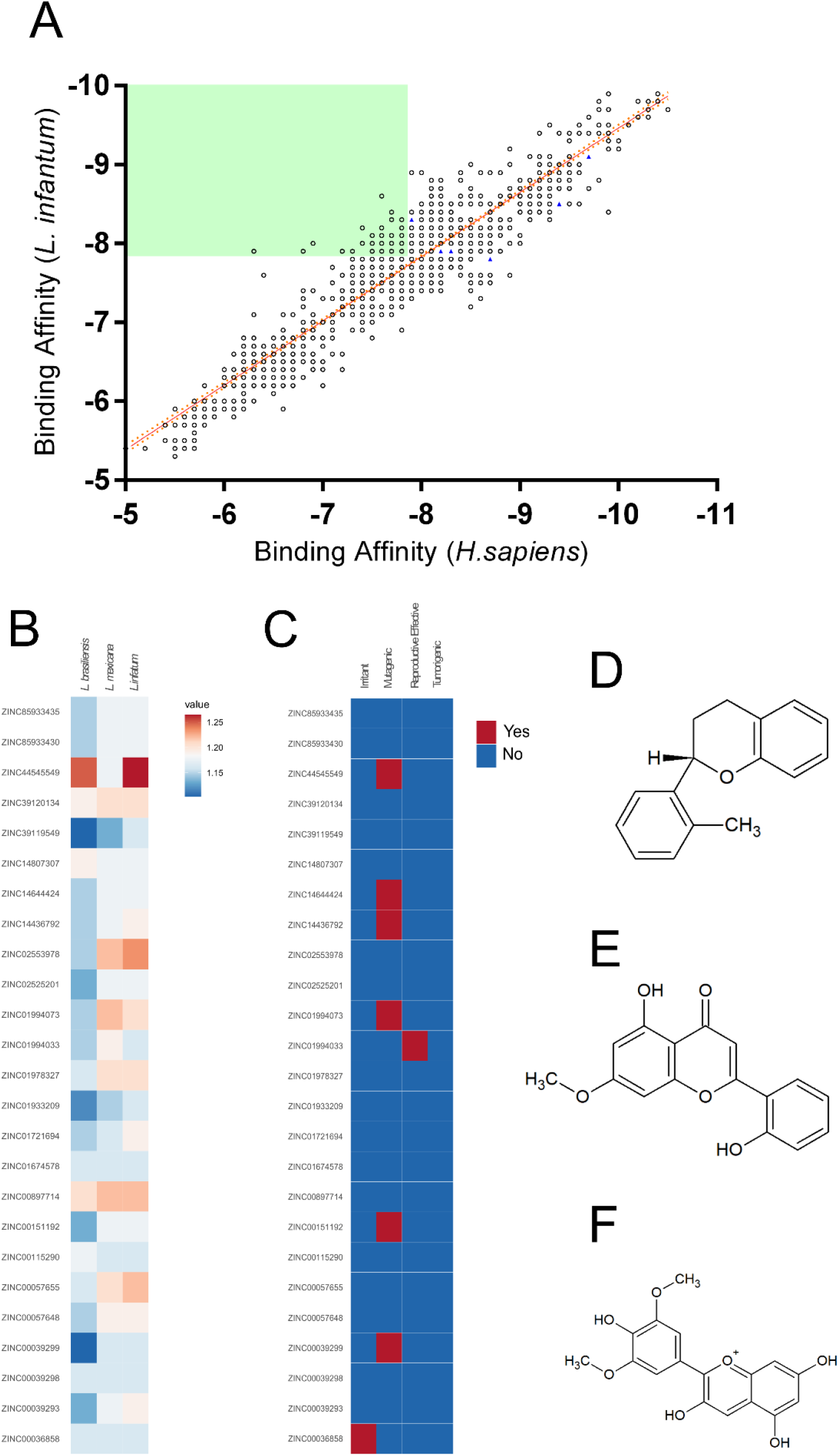
Virtual screening of the compounds selected from the NuBBE database. Binding affinities toward *L. infatum* and *H. sapiens* ARG targets were analyzed by linear regression and Pearson’s correlation coefficient. Solid orange line: linear regression; dotted orange lines: 95% confidence intervals. The solid green square was calculated using the maximum binding affinities of the 6 NPs (**A**). Normalized binding affinities heatmap of 25 selected compounds on *L. infatum, L. mexicana*, and *L. braziliensis* against their human homolog (**B**). Binary heatmap showing positive (red) or negative (blue) predicted toxicities (**C**). Chemical structure of ZINC39120134 (**D**), ZINC14807307 (**E**), and ZINC897714 (**F**).

### Molecular dynamics simulations (MDS)

*L. infantum* ARG is an enzyme with trimeric conformation (ChainA, ChainB, and ChainC) and its structure showed stable behavior during a 100 ns of MDS performed at pH 2 and pH 7 (Figure 2). Here we included the metal ions (Mn^+2^) and one hydroxyl molecule (OH^-1^) for each active site, and it was observed that some regions lose their structural conformation at pH 2.0 conditions (green color). Also, ARG at pH 2.0 shows substantial conformational changes concerning pH 7 (Figure 3A), and the high fluctuations in pH 2.0 allow water access (Figure 3B). In Figure 3C, the radius of gyration shows lower compaction of whole protein during the MDS at pH 7.0 than pH 2.0.

**Figure 2.**
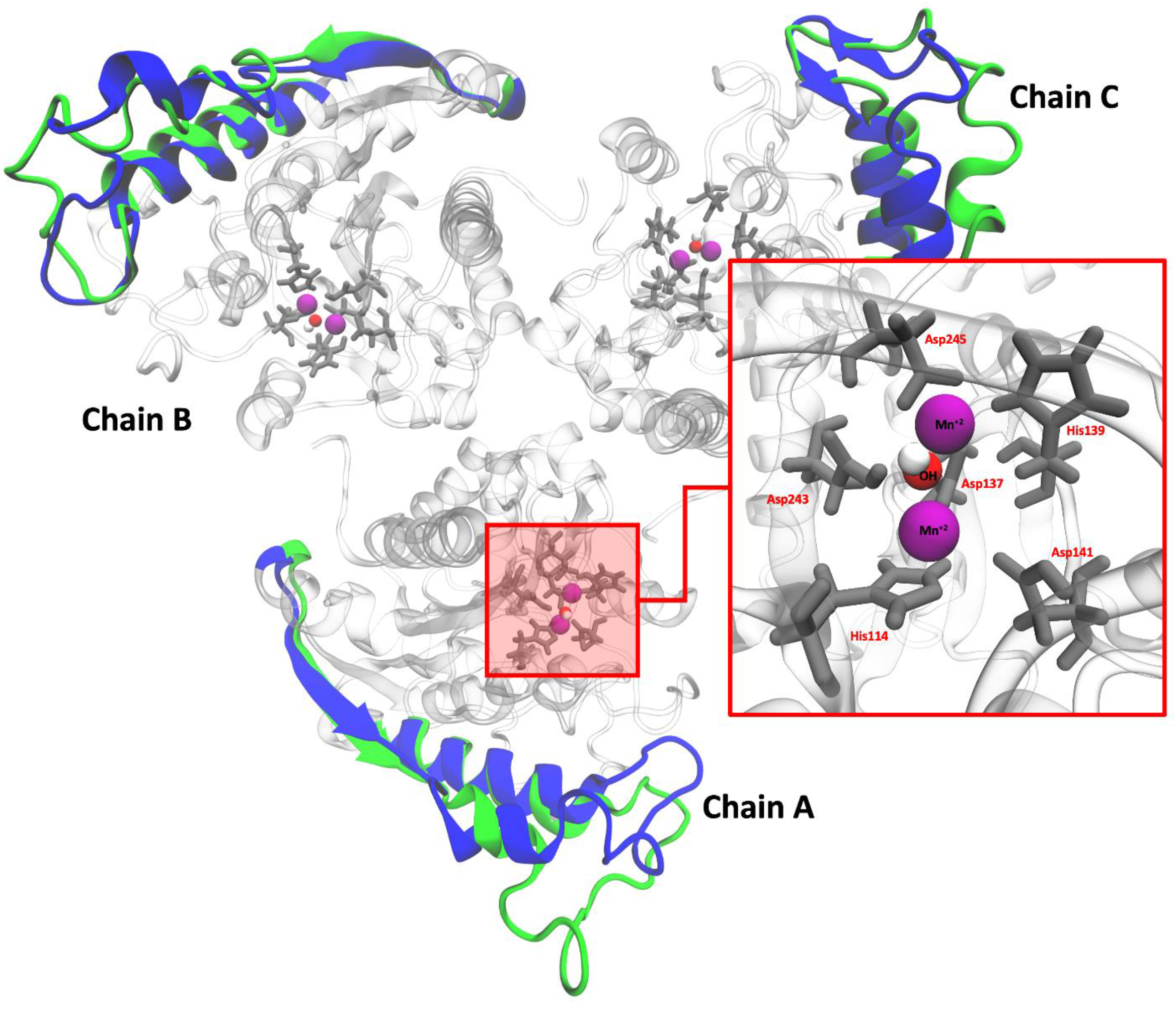
Structural conformation of Arginase with its active site. Color blue and green represent the cartoon representation of pH 2 and ely. The red box shows the active site of arginase.

**Figure 3.**
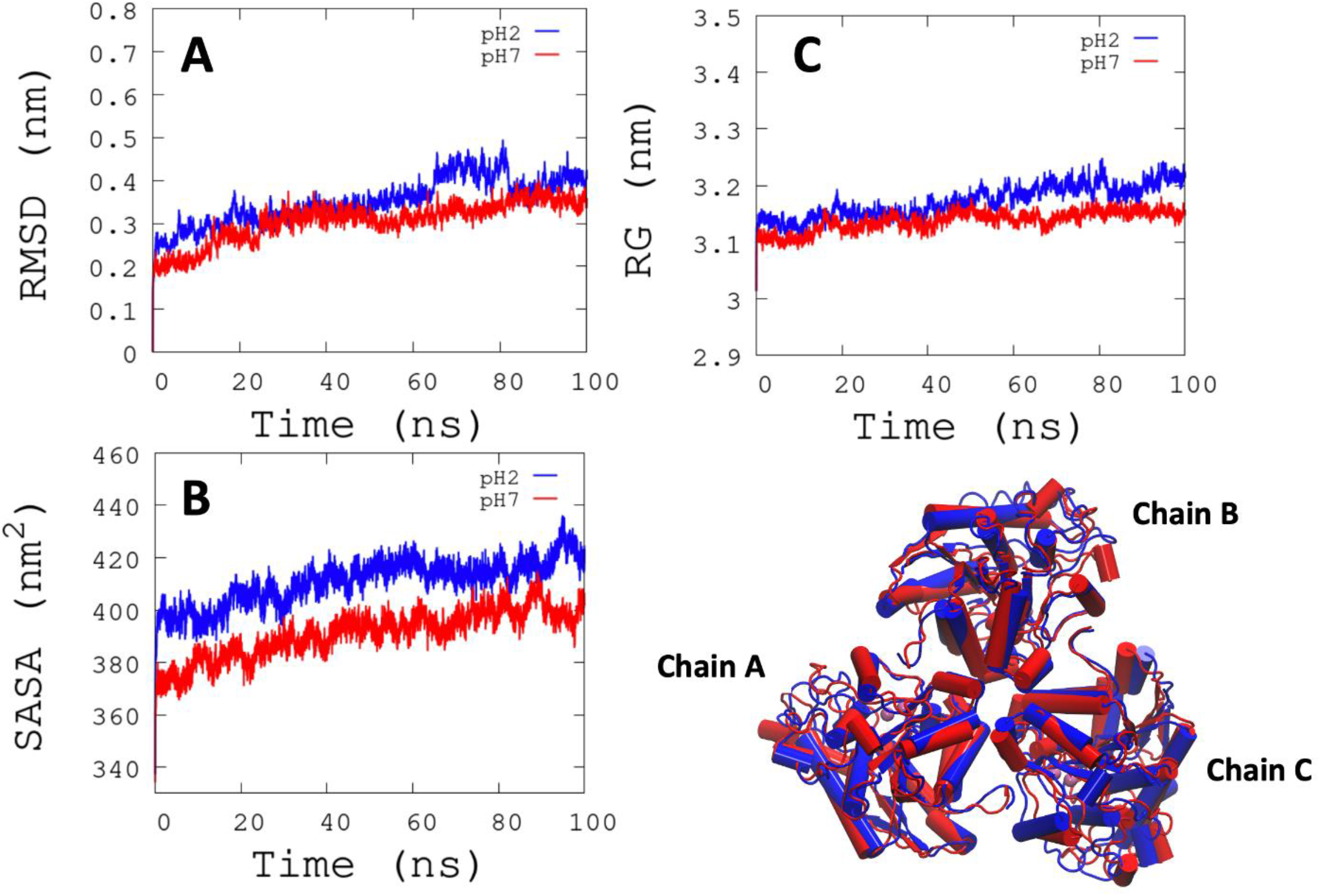
RMSD, SASA, and Radius of gyration analysis. A. RMSD is shown the conformational changes reported at pH 2. B. There is greater solvent access surface area to arginase at pH2. C. RG shows the same behavior as RMSD.

The report of the trajectory of each complex system (receptor-ligand) and the protein without ligand is shown in Figure 4. Since the Root-mean-squared deviation (RMSD) is a noteworthy analysis to verify the similarity between a protein-bound and not bound ligand [37]. The RMSD values in nm are presented that were taken from the A-chain of each protein in different pH conditions, whereas the receptor-ligand systems presented greater conformational changes in the substrate-binding site (Figure 4A). Likewise, Radius of Gyration (RG) analysis verifies the compactness of protein structures, whereas the lowest RG demonstrates the tightest packing and high conformational stability [38]. The results showed that at pH 2.0, low compactness and a large broadening of the macromolecule are reported (Figure 4B). Figure 4C shows the Root-mean-squared fluctuation (RMSF) *per residue* of the backbone, where high fluctuations were shown from residue 50 to 100 in both systems. From the receptor-ligand simulations’ results, we take each simulation’s last frames (Figure 5). The ligand ZINC897714 generates exciting interactions in the active center at the pH conditions evaluated and at pH 7.0, hydrogen bonds are observed, which benefits receptor-ligand coupling.

**Figure 4.**
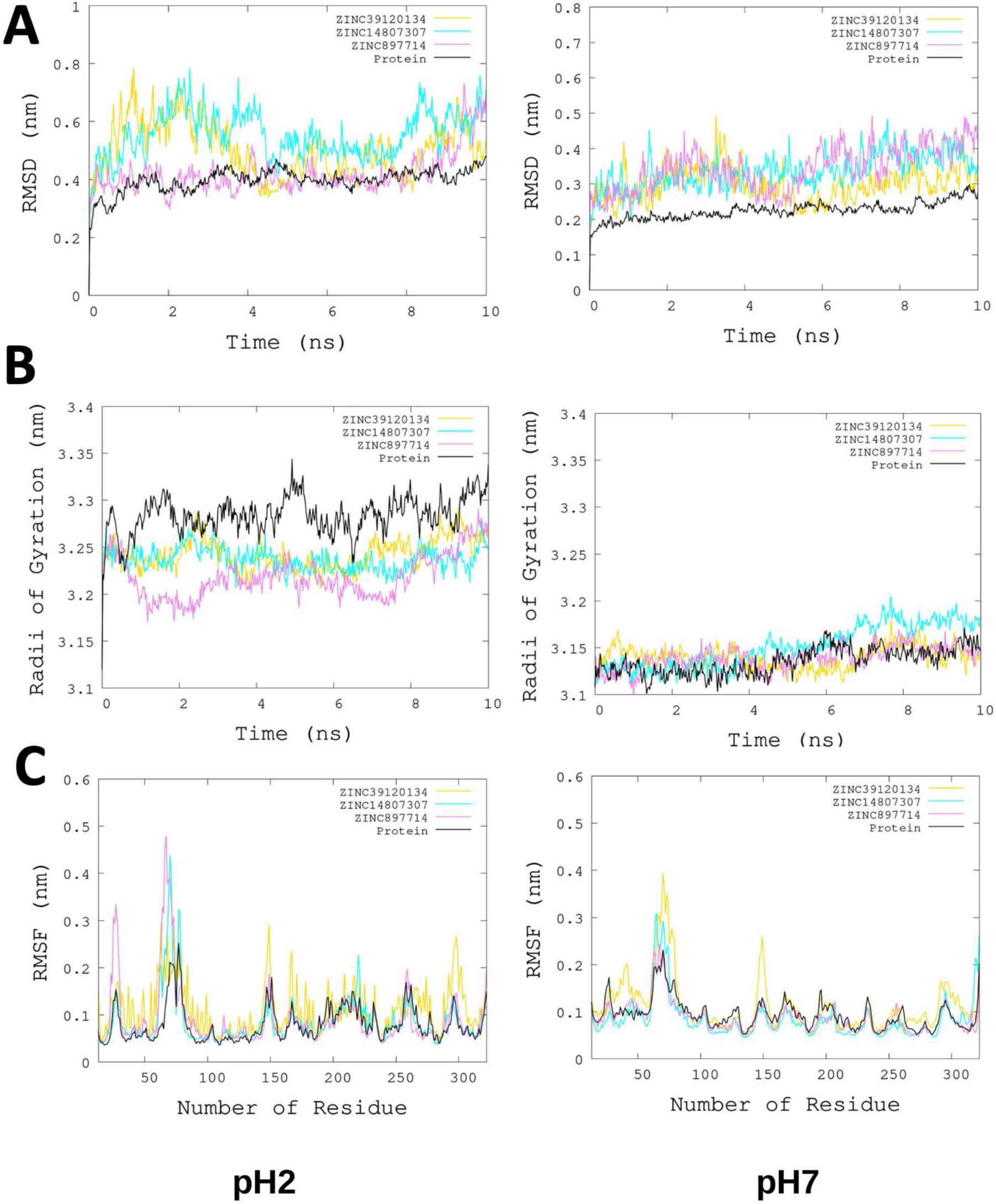
Plots of MD simulation of each complex. More significant conformational changes of arginase enzyme are shown at pH 2. A. RMSD plot of chain A concerning the whole protein. B. RG analysis. C. RMSF per residue of backbone.

**Figure 5.**
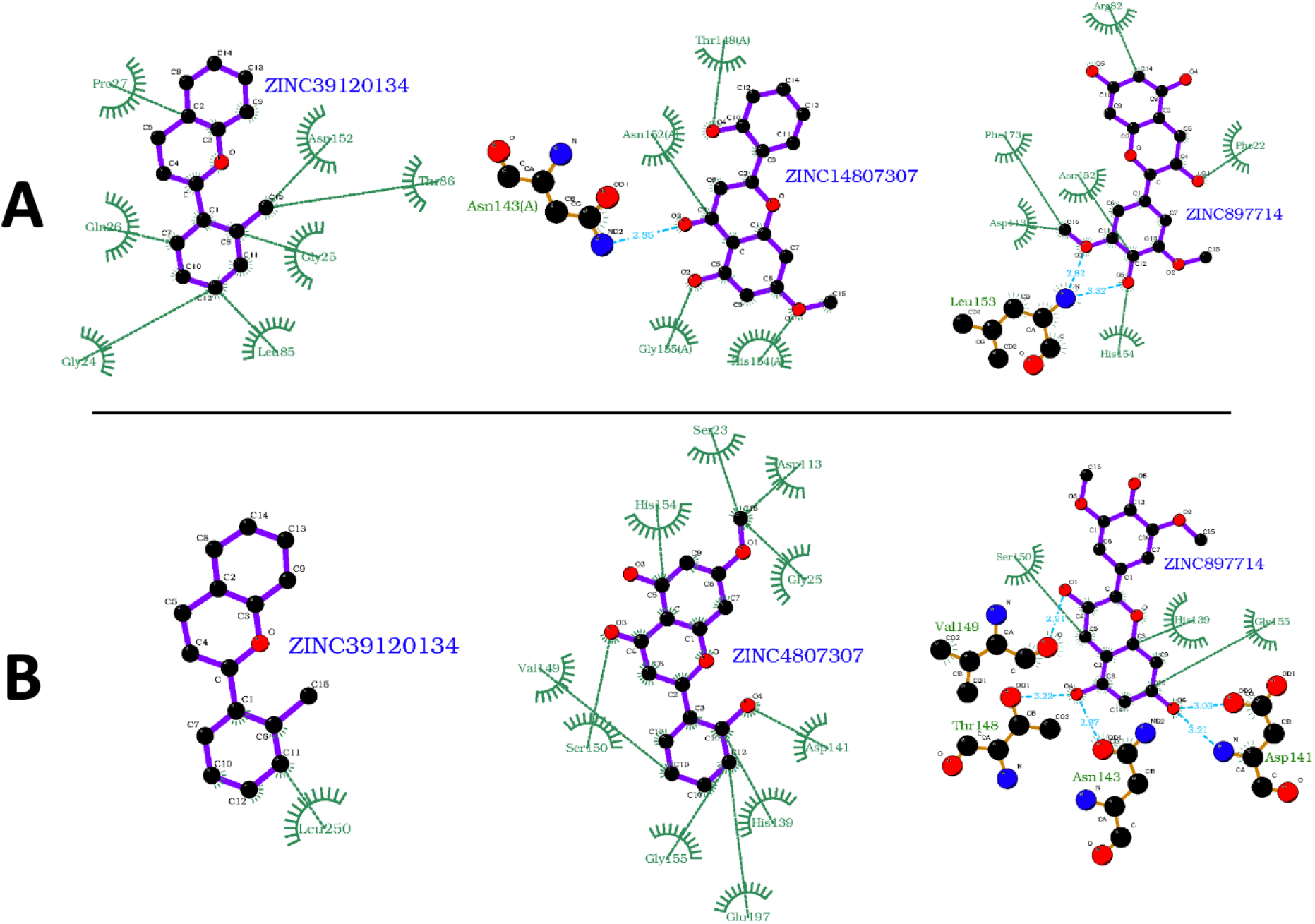
2D representation of the last frame of each complex. A. Last frame at pH 2. and B. Last frame at pH 7. Green represents the hydrophobic interaction between receptor-ligand, color sky blue represents the hydrogen bond interaction.

### MM-GBSA Analysis

The Binding Free Energy Analysis of pH 2.0 and pH 7.0 from the last 50 frames of each simulation is shown in Table 2. The propitious energetic contribution with a binding free energy of -23.68±2.76 kcal/mol (ZINC39120134/pH2) maximum and -13.45±2.58 kcal/mol (ZINC39120134/pH7) minimum were obtained. It is indicated that for the two pH conditions analyzed, the van der Waals energies (ΔEvdW) provided the highest energy contributions to the systems with ZINC39120134 and ZINC14807307. However, in the systems with ZINC897714, the electrostatic energies (ΔEele) contributed negatively to the binding receptor-ligand, which is attributed to the total net charge positive of the ligand and the pocket residues (induced by the protonation states at pH 2.0 and pH 7). Despite this, the solvation energies (ΔEgb) offset the positive electrostatic interactions, thus favorably contributing to the binding of ZINC897714 to ARG. These results show that the protonation states at a given pH can positively or negatively favor the enzyme-ligand binding, where it is expected that at pH above 7.0 the enzyme-ligand binding can be increased.

**Table 2.**
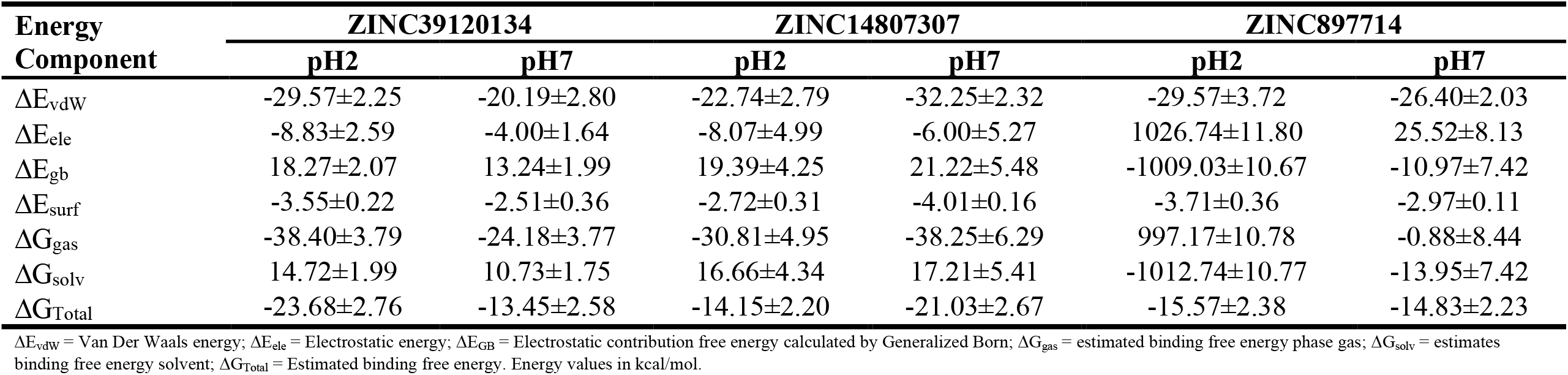
MM/GBSA binding free energy estimation

## Discussion

The World Health Organization (WHO) considers leishmaniasis to be one of the major neglected global diseases and responsible for millions of disability-adjusted life years (DALYs), representing one of the top burdens among the neglected tropical diseases [39]. Worldwide, 13 countries have a high burden of VL (Bangladesh, China, Ethiopia, Georgia, India, Kenya, Nepal, Paraguay, Somalia, South Sudan, Spain, Sudan, and Uganda), and 11 have a high burden of TL (Afghanistan, Algeria, Colombia, Iran, Morocco, Pakistan, Peru, Saudi Arabia, Syrian Arab Republic, Tunisia, and Turkey), while Brazil has a high burden of both clinical forms [40]. As VL is the most serious manifestation of the disease, any person with signs and symptoms and a confirmed diagnosis of VL warrants early systemic therapy, while TL treatment is based on the clinical presentation, host, and species type [41]. The range of currently available drugs for treating leishmaniasis is relatively small and it includes notably 62.5% of repurposed molecules, while less than 1% of all new drug candidates reached clinical trials in the last decades [42,43]. For these reasons, the investigation of new therapies has been very active recently, whereas a wide range of compounds has been identified as potential hits and leads [44].

A large component of biologically relevant chemical space is occupied by NPs; since they have a unique and vast chemical diversity [45] and this condition is no different when it comes to antileishmanial molecules, as several NPs such as alkaloids, coumarins, quinones, flavonoids, terpenoids, lignans, and neolignans have been described as active against various Leishmania species [46]. Some of these NPs molecules, that target *Leishmania* ARG have been studied for the development of new therapeutics; as Quercetin had shown antileishmanial activity *in vitro* [47–49] and *in vivo* [50,51]; Fisetin, Catechins, Resveratrol, and Cinnamic Acid Derivatives had significantly shown *in vitro* efficacy [30]; and, Epigallocatechin gallate [52], Gallic acid [53], and Rosmarinic acid [54]; showed promising *in vivo* activity. However, the translation of NPs to commercial drugs is limited by several factors, since the balance between biological activity and pharmaceutical properties related to chemical structure, such as pharmacokinetic profile, is a delicate matter [55,56].

*In silico*-based drug repositioning potential for discovering new applications for existing drugs and for developing new drugs in pharmaceutical research and the industry has gained importance [57,58]; whereas in the chemical structure and molecule information approach, the structural similarity is incorporated with molecular activity and other biological information to identify new associations [59]. The present work aimed to apply distinct bioinformatic tools to select analogs to NPs with known antileishmanial and anti-ARG activities; although results of the Quercetin analog, the anthocyanin Malvidin (ZINC897714; PubChem CID: 159287) showed favorable binding affinity to *L. infantum, L. mexicana*, and *L. braziliensis* ARG and no predicted toxicity. A major bottleneck of drug discovery on leishmaniasis was aimed at the *in-silico* workflow proposed, which is that compounds must show activity in the acidic environment of the phagolysosome [60]; thus, Malvidin showed stable receptor-ligand interaction and favorable binding free energy at pH 2 in MDS analysis. Also, the Target Product Profile (TPP) proposed by the Drugs for Neglected Diseases initiative (DNDi), includes regard for the oral route of administration for new candidates [61], while the ADME profile of the compound shows its potential for oral route administration and high bioavailability [62–64]. Furthermore, Malvidin has shown the potential to be an antioxidant, anti-hypertensive, anti-inflammatory, anti-obesity, anti-osteoarthritis, anti-proliferative, and anticancer drug candidate [65–69].

Anthocyanins are commonly found in many plants, while the most common types are cyanidin, delphinidin, pelargonidin, peonidin, petunidin, and malvidin; which are distributed in fruits and vegetables in 50%, 12%, 12%, 12%, 7%, and 7% proportion, respectively [70]. These molecules are more stable at a lower pH solution, and in such conditions, the flavylium cation formed enables the anthocyanin to be highly soluble in water [71]. The physicochemical properties offered by anthocyanins should be considered of interest for antileishmanial drug discovery since the parasite is adapted to live in parasitophorous vacuoles of infected macrophages in mammalian hosts, where it survives, proliferates, and is responsible for the development of the active disease [72]. Recently, the anthocyanidin profile of *Arrabidaea chica* has been examined and their antileishmanial activity was analyzed [73]; whereas Carajurin (PubChem CID: 44257040) showed the highest activity against the intracellular parasites, altering all parameters of in vitro infection [74]. Additionally, it has been shown that Carajurin leads to a decrease in the mitochondrial membrane potential, an increase in reactive oxygen species production, and cell death by late apoptosis in *L. amazonensis* [75].

Limitations of the present study should be also mentioned, such as the protein dynamics and complex stabilities with MDS lasting within nanoseconds scales (0–100 ns), while most structural dynamics and biological activities of proteins occur within timescales of microseconds and milliseconds [76]. However, it has been considered that MDS analyses of protein-ligand interactions and complex dynamics could be accurately informed by nanosecond timescales [77,78]. Also, the work did not include *in vitro* or *in vivo* validation of the results; however, it is worth noting that axenic form-based assays present disadvantages, such as the metabolic differences between the amastigote and promastigote stages that may lead to the selection of misleading candidates, which is verified by the lack of correlation between selected compounds in axenic forms screenings and intracellular amastigote assays [79]. Additionally, differential drug susceptibility and infectiveness among parasites isolated from patients who exhibited differences in clinical outcomes [80] and several biochemical pathways that are associated with drug resistance phenotype in *Leishmania* [81,82] highlight the importance of taking into consideration this knowledge when performing *in vitro* validation. Furthermore, many animal models are used for VL and TL drug candidates’ validation assays; however, their predictive validity is often low due to incomplete translation to the human disease. Besides that, for VL robust primary models, such as BALB/c mice and Syrian golden hamsters are widely used [83,84]; however, there are no validated animal models for TL since there are several species causing different clinical manifestations, which brings complexity to the validation of these models with features like humans concerning etiology, pathophysiology, symptomatology, and response to chemotherapeutics [85].

## Conclusion

In the first screening, this work identified three substances with natural products structural analogs with potential effects against *Leishmania* ARG using *in silico* analysis from the available data and research of natural products found in databases. The substances were: ZINC39120134 (3,4-dihydro-2-(2-methylphenyl)-(9CI)), ZINC14807307 (Echioidinin) and ZINC897714 (Malvidin), in which the most suitable compound was ZINC897714 showing favorable binding affinity to *L. infantum, L. mexicana*, and *L. braziliensis* ARG, no potential toxicity and stability at pH 2.0; important factors due to the acidic environment of the phagolysosomes of mammalian hosts. The ADME analysis predicted no toxic effect for Malvidin, showing it as a therapeutic possible for drug formulation. The RMSD simulations and predictions have demonstrated higher docking stabilities for the systems. The results presented in this work warrants further *in vitro* and *in vivo* studies using Malvidin to confirm its potential as a drug candidate against leishmaniasis.

## Methods

### Data collection

The search for natural products with antileishmanial and arginase activity, was performed at the Nuclei of Bioassays, Ecophysiology, and Biosynthesis of Natural Products Database (NuBBE_DB_) online web server (version 2017) (https://nubbe.iq.unesp.br/portal/nubbe-search.html, accessed on 23 January 2022), which contains the information of more than 2,000 natural products and derivatives [86]; while the “antileishmanial property” was selected in the biological properties segment of the web server. The bibliographic data extraction, regarding the compounds found in NuBBE_DB_, was performed from the National Center for Biotechnology Information (NCBI) databases (https://www.ncbi.nlm.nih.gov/pubmed/, accessed on 07 February 2022); and the simplified molecular-input line-entry system (SMILEs) was searched and retrieved from PubChem server (https://pubchem.ncbi.nlm.nih.gov/, accessed on 10 February 2022) [87]. Likewise, the physicochemical properties: Total Molecular Weight (MW), Octanol/Water Partition Coefficient (iLOGP), Number of H-Bond Acceptors (HBAs), Number of H-bond Donors (HBDs), and the Molecular Polar Surface Area (TPSA), for each compound were calculated within the Osiris DataWarrior v05.02.01 software [88]; and, the Rotatable Bonds (RB); Number of Heavy Atoms (NHA); and Synthetic accessibility (SynAcce) were calculated within SwissADME server (http://www.swissadme.ch/index.php, accessed on 15 February 2022) [89].

### Structural analogs search and virtual screening

The SMILES from the compounds were used for high throughput screening to investigate structural analogs by the SwissSimilarity server (http://www.swisssimilarity.ch/index.php, accessed on 01 March 2022) [90]; whereas the commercial class of compounds was selected and the Zinc-drug like compound library with the combined screening method were chosen for the high throughput screening (HTS) to achieve the best structural analogs. Threshold values for positivity were selected by default parameters.

Also, the FASTA sequences of the ARG sequences from *Leishmania infantum* (A4IB49), *L. mexicana* (Q6TUJ5), *L. brasiliensis* (A4HMH0), and *Homo sapiens* (P05089) were retrieved from UniProt database (http://www.uniprot.org/), accessed on 03 March 2022), and subjected to automated modeling in SWISS-MODEL [91]. Furthermore, the compounds were imported into OpenBabel within the Python Prescription Virtual Screening Tool [92] and subjected to energy minimization. PyRx performs structure-based virtual screening applying docking simulations using the AutoDock Vina tool [93], whereas the drug targets were uploaded as macromolecules. For the analysis, the search space encompassed the whole of the modeled 3D models; and the docking simulation was then run at an exhaustiveness of 8 and set to only output the lowest energy pose. The Osiris DataWarrior software was employed to calculate the potential tumorigenic, mutagenic, reproductive effect, and irritant action of selected compounds predicted by comparison to a precompiled fragment library derived from the RTECS (Registry of Toxic Effects of Chemical Substances) database [88].

### Molecular dynamics simulation

Ligands preparation was based on the results from the virtual screening analysis; while the geometry optimization of these compounds was made in the Avogadro v. 1.2.0 program [94] and ACPYPE [95] was employed to generate the topologies and parameters for molecular dynamics (MD) simulation. We determined the 3D structural conformation of *L. infantum* ARG by homology modeling with *L. mexicana* arginase (PDB ID: 4ITY) as a template in the SwissModel online server [91] and afterward we determined the protonation/deprotonation states at pH 2.0 and pH 7.0 in the PDB2PQR [96]. Since ARG is a trimeric metalloprotein with three active sites binding to two manganese atoms (Mn^+2^), we fixed the Mn^+2^ coordination with active site residues and a hydroxyl molecule (OH^-1^), considering the following coordination: first MN^+2^ with His114 (ND1), ASP137 (OD2), ASP141 (OD2), ASP243 (OD2) and the second MN^+2^ with ASP137 (OD1), HIS139 (ND1), ASP243 (OD1) and ASP245 (OD2). The MD simulation was reproduced in GROMACS v. 2020 [97], considering the AMBER99 [98] force field. The systems were solvated with the TIP3P water model, and Na^+1^ or Cl-^1^ ions were added for neutralization. The box size was 12×12×12. Thus, the energy minimization was performed with the steep-descent algorithm with 20000 steps of calculation. The MD simulation was done in two steps; the first step was in the canonical ensemble NVT considering distance restraint of Mn^+2^ to the active site by 5ns. The second step was the MD production in the isothermal-isobaric ensemble NPT with a time of 100ns. The V-rescale [99] thermostat was used to regulate the temperature at 309.65 K and the Parrinello-Rahman barostat at a reference pressure of 1 bar. Molecular docking was done with the DockThor online server [100]; in the last frame, the molecular docking at two pH conditions was used as a receptor. A grid was considered in the active site of arginase (chain A). The complex models with the best scores were chosen, and these were subsequently simulated in the isothermal-isobaric ensemble NPT for 10 ns. Gibbs free energy was calculated by the Molecular Mechanics-Generalized Born Surface Area (MM-GBSA) [101] method in gmx_MMPBSA X tool based on AMBER’s MMPBSA.py, and AmberTools20 [102] package was used.

### Statistical analysis

Results were entered into Microsoft Excel (version 10.0, Microsoft Corporation, Redmond, WA, USA) spreadsheets and analyzed by GraphPad Prism version 9.4.0 (673) for Windows, GraphPad Software, San Diego, California USA, “www.graphpad.com”. To evaluate the correlation between the binding affinities of the compounds against the protein targets were placed in a linear regression plot and analyzed by Pearson’s correlation coefficient, differences were considered significant when *p*< 0.05. Heatmaps were constructed in the R programming environment (version 4.0.3) using the “*heatmap* 2” function in the package “*gplots*” [103].

## Author Contributions

Conceptualization: H.L.B.-C. and M.A.C.-F.; data curation: L.D.G.-M., M.A.C.-P, C.S.F., G.S.V.T., D.L.P, and E.A.F.C; formal analysis: H.L.B.-C. and M.A.C.-F.; funding acquisition: H.L.B.-C. and M.A.C.-F.; investigation: H.L.B.-C., L.D.G.-M., D.L.P., G.S.V.T., and C.S.F.; methodology: H.L.B.-C. and M.A.C.-F.; writing—review & editing: H.L.B.-C., E.A.F.C., and M.A.C.-F. All authors have read and agreed to the published version of the manuscript.

## Institutional Review Board Statement

Not applicable.

## Informed Consent Statement

Not applicable.

## Data Availability Statement

Not applicable.

## Acknowledgments

Not applicable.

## References

1. Tuon, F.F.; Amato Neto, V.; Sabbaga Amato, V. Leishmania : Origin, Evolution and Future since the Precambrian. FEMS Immunology & Medical Microbiology 2008, 54, 158–166, doi:10.1111/j.1574-695X.2008.00455.x.

2. Alvar, J.; Vélez, I.D.; Bern, C.; Herrero, M.; Desjeux, P.; Cano, J.; Jannin, J.; Boer, M. den Leishmaniasis Worldwide and Global Estimates of Its Incidence. PLoS ONE 2012, 7, e35671, doi:10.1371/journal.pone.0035671.

3. Hotez, P.J. The Rise of Leishmaniasis in the Twenty-First Century. Transactions of The Royal Society of Tropical Medicine and Hygiene 2018, 112, 421–422, doi:10.1093/trstmh/try075.

4. Oryan, A.; Akbari, M. Worldwide Risk Factors in Leishmaniasis. Asian Pacific Journal of Tropical Medicine 2016, 9, 925–932, doi:10.1016/j.apjtm.2016.06.021.

5. Curtin, J.M.; Aronson, N.E. Leishmaniasis in the United States: Emerging Issues in a Region of Low Endemicity. Microorganisms 2021, 9, 578, doi:10.3390/microorganisms9030578.

6. Ready, P.D. Leishmaniasis Emergence in Europe. Eurosurveillance 2010, 15, doi:10.2807/ese.15.10.19505-en.

7. Lauletta Lindoso, J.A.; Alves Cunha, M.; Queiroz, I.; Valente Moreira, C.H. Leishmaniasis–HIV Coinfection: Current Challenges. HIV/AIDS - Research and Palliative Care 2016, Volume 8, 147–156, doi:10.2147/HIV.S93789.

8. Dostálová, A.; Volf, P. Leishmania Development in Sand Flies: Parasite-Vector Interactions Overview. Parasit Vectors 2012, 5, 276, doi:10.1186/1756-3305-5-276.

9. Akhoundi, M.; Kuhls, K.; Cannet, A.; Votýpka, J.; Marty, P.; Delaunay, P.; Sereno, D. A Historical Overview of the Classification, Evolution, and Dispersion of Leishmania Parasites and Sandflies. PLOS Neglected Tropical Diseases 2016, 10, e0004349, doi:10.1371/journal.pntd.0004349.

10. Kaye, P.; Scott, P. Leishmaniasis: Complexity at the Host–Pathogen Interface. Nature Reviews Microbiology 2011, 9, 604–615, doi:10.1038/nrmicro2608.

11. Burza, S.; Croft, S.L.; Boelaert, M. Leishmaniasis. The Lancet 2018, 392, 951–970, doi:10.1016/S0140-6736(18)31204-2.

12. Kaye, P.M.; Mohan, S.; Mantel, C.; Malhame, M.; Revill, P.; le Rutte, E.; Parkash, V.; Layton, A.M.; Lacey, C.J.N.; Malvolti, S. Overcoming Roadblocks in the Development of Vaccines for Leishmaniasis. Expert Review of Vaccines 2021, 20, 1419–1430, doi:10.1080/14760584.2021.1990043.

13. Lindoso, J.A.L.; Costa, J.M.L.; Queiroz, I.T.; Goto, H. Review of the Current Treatments for Leishmaniases. Research and Reports in Tropical Medicine 2012, 3, 69–77, doi:10.2147/RRTM.S24764.

14. Chakravarty, J.; Sundar, S. Current and Emerging Medications for the Treatment of Leishmaniasis. Expert Opinion on Pharmacotherapy 2019, 20, 1251–1265, doi:10.1080/14656566.2019.1609940.

15. Newman, D.J.; Cragg, G.M. Natural Products as Sources of New Drugs over the Nearly Four Decades from 01/1981 to 09/2019. Journal of Natural Products 2020, 83, 770–803, doi:10.1021/acs.jnatprod.9b01285.

16. Ribeiro, T.G.; Nascimento, A.M.; Henriques, B.O.; Chávez-Fumagalli, M. a.; Franca, J.R.; Duarte, M.C.; Lage, P.S.; Andrade, P.H.R.; Lage, D.P.; Rodrigues, L.B.; et al. Antileishmanial Activity of Standardized Fractions of Stryphnodendron Obovatum (Barbatimão) Extract and Constituent Compounds. Journal of Ethnopharmacology 2015, 165, 238–242, doi:10.1016/j.jep.2015.02.047.

17. Lage, P.S.; de Andrade, P.H.R.; Lopes, A.D.S.; Chávez Fumagalli, M.A.; Valadares, D.G.; Duarte, M.C.; Pagliara Lage, D.; Costa, L.E.; Martins, V.T.; Ribeiro, T.G.; et al. Strychnos Pseudoquina and Its Purified Compounds Present an Effective In Vitro Antileishmanial Activity. Evidence-Based Complementary and Alternative Medicine 2013, 2013, 1–9, doi:10.1155/2013/304354.

18. Ribeiro, T.G.; Chávez-Fumagalli, M. a.; Valadares, D.G.; Franca, J.R.; Lage, P.S.; Duarte, M.C.; Andrade, P.H.R.; Martins, V.T.; Costa, L.E.; Arruda, A.L.A.; et al. Antileishmanial Activity and Cytotoxicity of Brazilian Plants. Experimental Parasitology 2014, 143, 60–68, doi:10.1016/j.exppara.2014.05.004.

19. Lage, P.S.; Chávez-Fumagalli, M. a.; Mesquita, J.T.; Mata, L.M.; Fernandes, S.O. a.; Cardoso, V.N.; Soto, M.; Tavares, C. a. P.; Leite, J.P. v.; Tempone, A.G.; et al. Antileishmanial Activity and Evaluation of the Mechanism of Action of Strychnobiflavone Flavonoid Isolated from Strychnos Pseudoquina against Leishmania Infantum. Parasitology Research 2015, 114, 4625–4635, doi:10.1007/s00436-015-4708-4.

20. Goyzueta-Mamani, L.D.; Barazorda-Ccahuana, H.L.; Mena-Ulecia, K.; Chávez-Fumagalli, M.A. Antiviral Activity of Metabolites from Peruvian Plants against SARS-CoV-2: An In Silico Approach. Molecules 2021, 26, 3882, doi:10.3390/molecules26133882.

21. Goyzueta-Mamani, L.D.; Barazorda-Ccahuana, H.L.; Chávez-Fumagalli, M.A. F. Alvarez, K.L.; Aguilar-Pineda, J.A.; Vera-Lopez, K.J.; Lino Cardenas, C.L. In Silico Analysis of Metabolites from Peruvian Native Plants as Potential Therapeutics against Alzheimer’s Disease. Molecules 2022, 27, 918, doi:10.3390/molecules27030918.

22. Durieu, E.; Prina, E.; Leclercq, O.; Oumata, N.; Gaboriaud-Kolar, N.; Vougogiannopoulou, K.; Aulner, N.; Defontaine, A.; No, J.H.; Ruchaud, S.; et al. From Drug Screening to Target Deconvolution: A Target-Based Drug Discovery Pipeline Using Leishmania Casein Kinase 1 Isoform 2 To Identify Compounds with Antileishmanial Activity. Antimicrobial Agents and Chemotherapy 2016, 60, 2822–2833, doi:10.1128/AAC.00021-16.

23. Bell, A.S.; Mills, J.E.; Williams, G.P.; Brannigan, J.A.; Wilkinson, A.J.; Parkinson, T.; Leatherbarrow, R.J.; Tate, E.W.; Holder, A.A.; Smith, D.F. Selective Inhibitors of Protozoan Protein N-Myristoyltransferases as Starting Points for Tropical Disease Medicinal Chemistry Programs. PLoS Neglected Tropical Diseases 2012, 6, e1625, doi:10.1371/journal.pntd.0001625.

24. Chávez-Fumagalli, M.A.; Lage, D.P.; Tavares, G.S.V.; Mendonça, D.V.C.; Dias, D.S.; Ribeiro, P.A.F.; Ludolf, F.; Costa, L.E.; Coelho, V.T.S.; Coelho, E.A.F. In Silico Leishmania Proteome Mining Applied to Identify Drug Target Potential to Be Used to Treat against Visceral and Tegumentary Leishmaniasis. Journal of Molecular Graphics and Modelling 2019, 87, 89–97, doi:10.1016/j.jmgm.2018.11.014.

25. Raj, S.; Sasidharan, S.; Balaji, S.N.; Saudagar, P. An Overview of Biochemically Characterized Drug Targets in Metabolic Pathways of Leishmania Parasite. Parasitology Research 2020, 119, doi:10.1007/s00436-020-06736-x.

26. Roberts, S.C.; Tancer, M.J.; Polinsky, M.R.; Michael Gibson, K.; Heby, O.; Ullman, B. Arginase Plays a Pivotal Role in Polyamine Precursor Metabolism in Leishmania: Characterization of Gene Deletion Mutants. Journal of Biological Chemistry 2004, 279, doi:10.1074/jbc.M402042200.

27. Pessenda, G.; Silva, J.S. Arginase and Its Mechanisms in Leishmania Persistence. Parasite Immunology 2020, 42, doi:10.1111/pim.12722.

28. da Silva, E.R.; Maquiaveli, C. do C.; Magalhães, P.P. The Leishmanicidal Flavonols Quercetin and Quercitrin Target Leishmania (Leishmania) Amazonensis Arginase. Experimental Parasitology 2012, 130, 183–188, doi:10.1016/j.exppara.2012.01.015.

29. Wulsten, I.F.; Costa-Silva, T.A.; Mesquita, J.T.; Lima, M.L.; Galuppo, M.K.; Taniwaki, N.N.; Borborema, S.E.T.; da Costa, F.B.; Schmidt, T.J.; Tempone, A.G. Investigation of the Anti-Leishmania (Leishmania) Infantum Activity of Some Natural Sesquiterpene Lactones. Molecules 2017, 22, doi:10.3390/molecules22050685.

30. Carter, N.S.; Stamper, B.D.; Elbarbry, F.; Nguyen, V.; Lopez, S.; Kawasaki, Y.; Poormohamadian, R.; Roberts, S.C. Natural Products That Target the Arginase in Leishmania Parasites Hold Therapeutic Promise. Microorganisms 2021, 9, 267, doi:10.3390/microorganisms9020267.

31. de Sousa, L.R.F.; Ramalho, S.D.; Burger, M.C. de M.; Nebo, L.; Fernandes, J.B.; da Silva, M.F. das G.F.; Iemma, M.R. da C.; Corrêa, C.J.; Souza, D.H.F. de; Lima, M.I.S.; et al. Isolation of Arginase Inhibitors from the Bioactivity-Guided Fractionation of Byrsonima Coccolobifolia Leaves and Stems. Journal of Natural Products 2014, 77, 392–396, doi:10.1021/np400717m.

32. Chagas, C.M.; Moss, S.; Alisaraie, L. Drug Metabolites and Their Effects on the Development of Adverse Reactions: Revisiting Lipinski’s Rule of Five. International Journal of Pharmaceutics 2018, 549, 133–149, doi:10.1016/j.ijpharm.2018.07.046.

33. Bickerton, G.R.; Paolini, G. v.; Besnard, J.; Muresan, S.; Hopkins, A.L. Quantifying the Chemical Beauty of Drugs. Nature Chemistry 2012, 4, 90–98, doi:10.1038/nchem.1243.

34. Ertl, P.; Schuffenhauer, A. Estimation of Synthetic Accessibility Score of Drug-like Molecules Based on Molecular Complexity and Fragment Contributions. Journal of Cheminformatics 2009, 1, 8, doi:10.1186/1758-2946-1-8.

35. Fernández, O.L.; Diaz-Toro, Y.; Ovalle, C.; Valderrama, L.; Muvdi, S.; Rodríguez, I.; Gomez, M.A.; Saravia, N.G. Miltefosine and Antimonial Drug Susceptibility of Leishmania Viannia Species and Populations in Regions of High Transmission in Colombia. PLoS Neglected Tropical Diseases 2014, 8, e2871, doi:10.1371/journal.pntd.0002871.

36. Baek, K.-H.; Piel, L.; Rosazza, T.; Prina, E.; Späth, G.F.; No, J.H. Infectivity and Drug Susceptibility Profiling of Different Leishmania-Host Cell Combinations. Pathogens 2020, 9, 393, doi:10.3390/pathogens9050393.

37. Coutsias, E.A.; Wester, M.J. RMSD and Symmetry. Journal of Computational Chemistry 2019, 40, 1496–1508, doi:10.1002/jcc.25802.

38. Lobanov, M.Yu.; Bogatyreva, N.S.; Galzitskaya, O. v. Radius of Gyration as an Indicator of Protein Structure Compactness. Molecular Biology 2008, 42, 623–628, doi:10.1134/S0026893308040195.

39. Vos, T.; Lim, S.S.; Abbafati, C.; Abbas, K.M.; Abbasi, M.; Abbasifard, M.; Abbasi-Kangevari, M.; Abbastabar, H.; Abd-Allah, F.; Abdelalim, A.; et al. Global Burden of 369 Diseases and Injuries in 204 Countries and Territories, 1990–2019: A Systematic Analysis for the Global Burden of Disease Study 2019. The Lancet 2020, 396, 1204–1222, doi:10.1016/S0140-6736(20)30925-9.

40. Torres-Guerrero, E.; Quintanilla-Cedillo, M.R.; Ruiz-Esmenjaud, J.; Arenas, R. Leishmaniasis: A Review. F1000Res 2017, 6, 750, doi:10.12688/f1000research.11120.1.

41. Mann, S.; Frasca, K.; Scherrer, S.; Henao-Martínez, A.F.; Newman, S.; Ramanan, P.; Suarez, J.A. A Review of Leishmaniasis: Current Knowledge and Future Directions. Current Tropical Medicine Reports 2021, 8, 121–132, doi:10.1007/s40475-021-00232-7.

42. Braga, S.S. Multi-Target Drugs Active against Leishmaniasis: A Paradigm of Drug Repurposing. European Journal of Medicinal Chemistry 2019, 183, 111660, doi:10.1016/j.ejmech.2019.111660.

43. de Rycker, M.; Baragaña, B.; Duce, S.L.; Gilbert, I.H. Challenges and Recent Progress in Drug Discovery for Tropical Diseases. Nature 2018, 559.

44. Olías-Molero, A.I.; de la Fuente, C.; Cuquerella, M.; Torrado, J.J.; Alunda, J.M. Antileishmanial Drug Discovery and Development: Time to Reset the Model? Microorganisms 2021, 9, 2500, doi:10.3390/microorganisms9122500.

45. Rosén, J.; Gottfries, J.; Muresan, S.; Backlund, A.; Oprea, T.I. Novel Chemical Space Exploration via Natural Products. Journal of Medicinal Chemistry 2009, 52, 1953–1962, doi:10.1021/jm801514w.

46. Gervazoni, L.F.O.; Barcellos, G.B.; Ferreira-Paes, T.; Almeida-Amaral, E.E. Use of Natural Products in Leishmaniasis Chemotherapy: An Overview. Frontiers in Chemistry 2020, 8, doi:10.3389/fchem.2020.579891.

47. Fonseca-Silva, F.; Inacio, J.D.F.; Canto-Cavalheiro, M.M.; Almeida-Amaral, E.E. Reactive Oxygen Species Production by Quercetin Causes the Death of Leishmania Amazonensis Intracellular Amastigotes. Journal of Natural Products 2013, 76, doi:10.1021/np400193m.

48. Vila-Nova, N.S.; Morais, S.M.; Falcão, M.J.C.; Bevilaqua, C.M.L.; Rondon, F.C.M.; Wilson, M.E.; Vieira, I.G.P.; Andrade, H.F. Leishmanicidal and Cholinesterase Inhibiting Activities of Phenolic Compounds of Dimorphandra Gardneriana and Platymiscium Floribundum, Native Plants from Caatinga Biome. Pesquisa Veterinaria Brasileira 2012, 32, doi:10.1590/S0100-736X2012001100015.

49. Cataneo, A.H.D.; Tomiotto-Pellissier, F.; Miranda-Sapla, M.M.; Assolini, J.P.; Panis, C.; Kian, D.; Yamauchi, L.M.; Colado Simão, A.N.; Casagrande, R.; Pinge-Filho, P.; et al. Quercetin Promotes Antipromastigote Effect by Increasing the ROS Production and Anti-Amastigote by Upregulating Nrf2/HO-1 Expression, Affecting Iron Availability. Biomedicine & Pharmacotherapy 2019, 113, 108745, doi:10.1016/j.biopha.2019.108745.

50. Sousa-Batista, A.J.; Poletto, F.S.; Philipon, C.I.M.S.; Guterres, S.S.; Pohlmann, A.R.; Rossi-Bergmann, B. Lipid-Core Nanocapsules Increase the Oral Efficacy of Quercetin in Cutaneous Leishmaniasis. Parasitology 2017, 144, doi:10.1017/S003118201700097X.

51. Tasdemir, D.; Kaiser, M.; Brun, R.; Yardley, V.; Schmidt, T.J.; Tosun, F.; Rüedi, P. Antitrypanosomal and Antileishmanial Activities of Flavonoids and Their Analogues: In Vitro, in Vivo, Structure-Activity Relationship, and Quantitative Structure-Activity Relationship Studies. Antimicrobial Agents and Chemotherapy 2006, 50, doi:10.1128/AAC.50.4.1352-1364.2006.

52. Sosa, A.M.; Álvarez, A.M.; Bracamonte, E.; Korenaga, M.; Marco, J.D.; Barroso, P.A. Efficacy of Topical Treatment with (-)-Epigallocatechin Gallate, a Green Tea Catechin, in Mice with Cutaneous Leishmaniasis. Molecules 2020, 25, doi:10.3390/molecules25071741.

53. de Moraes Alves, M.M.; Arcanjo, D.D.R.; Figueiredo, K.A.; de Sousa Macêdo Oliveira, J.S.; Viana, F.J.C.; de Sousa Coelho, E.; Lopes, G.L.N.; Gonçalves, J.C.R.; Carvalho, A.L.M.; dos Santos Rizzo, M.; et al. Gallic and Ellagic Acids Are Promising Adjuvants to Conventional Amphotericin B for the Treatment of Cutaneous Leishmaniasis. Antimicrobial Agents and Chemotherapy 2020, 64, doi:10.1128/AAC.00807-20.

54. Montrieux, E.; Perera, W.H.; García, M.; Maes, L.; Cos, P.; Monzote, L. In Vitro and in Vivo Activity of Major Constituents from Pluchea Carolinensis against Leishmania Amazonensis. Parasitology Research 2014, 113, 2925–2932, doi:10.1007/s00436-014-3954-1.

55. Atanasov, A.G.; Zotchev, S.B.; Dirsch, V.M.; Supuran, C.T. Natural Products in Drug Discovery: Advances and Opportunities. Nature Reviews Drug Discovery 2021, 20, 200–216, doi:10.1038/s41573-020-00114-z.

56. Batiha, G.E.-S.; Beshbishy, A.M.; Ikram, M.; Mulla, Z.S.; El-Hack, M.E.A.; Taha, A.E.; Algammal, A.M.; Elewa, Y.H.A. The Pharmacological Activity, Biochemical Properties, and Pharmacokinetics of the Major Natural Polyphenolic Flavonoid: Quercetin. Foods 2020, 9, 374, doi:10.3390/foods9030374.

57. Pushpakom, S.; Iorio, F.; Eyers, P.A.; Escott, K.J.; Hopper, S.; Wells, A.; Doig, A.; Guilliams, T.; Latimer, J.; McNamee, C.; et al. Drug Repurposing: Progress, Challenges and Recommendations. Nature Reviews Drug Discovery 2019, 18, 41–58, doi:10.1038/nrd.2018.168.

58. Mullins, J.G.L. Drug Repurposing in Silico Screening Platforms. Biochemical Society Transactions 2022, 50, 747–758, doi:10.1042/BST20200967.

59. Jarada, T.N.; Rokne, J.G.; Alhajj, R. A Review of Computational Drug Repositioning: Strategies, Approaches, Opportunities, Challenges, and Directions. Journal of Cheminformatics 2020, 12, 46, doi:10.1186/s13321-020-00450-7.

60. Alcântara, L.M.; Ferreira, T.C.S.; Gadelha, F.R.; Miguel, D.C. Challenges in Drug Discovery Targeting TriTryp Diseases with an Emphasis on Leishmaniasis. International Journal for Parasitology: Drugs and Drug Resistance 2018, 8, 430–439, doi:10.1016/j.ijpddr.2018.09.006.

61. Uliana, S.R.B.; Trinconi, C.T.; Coelho, A.C. Chemotherapy of Leishmaniasis: Present Challenges. Parasitology 2018, 145, 464–480, doi:10.1017/S0031182016002523.

62. Eker, M.E.; Aaby, K.; Budic-Leto, I.; Brncic, S.R.; El, S.N.; Karakaya, S.; Simsek, S.; Manach, C.; Wiczkowski, W.; de Pascual-Teresa, S. A Review of Factors Affecting Anthocyanin Bioavailability: Possible Implications for the Inter-Individual Variability. Foods 2020, 9, doi:10.3390/foods9010002.

63. Fagundes, F.L.; Pereira, Q.C.; Zarricueta, M.L.; dos Santos, R. de C. Malvidin Protects against and Repairs Peptic Ulcers in Mice by Alleviating Oxidative Stress and Inflammation. Nutrients 2021, 13, doi:10.3390/nu13103312.

64. Bub, A.; Watzl, B.; Heeb, D.; Rechkemmer, G.; Briviba, K. Malvidin-3-Glucoside Bioavailability in Humans after Ingestion of Red Wine, Dealcoholized Red Wine and Red Grape Juice. European Journal of Nutrition 2001, 40, doi:10.1007/s003940170011.

65. Ma, Y.; Li, Y.; Zhang, H.; Wang, Y.; Wu, C.; Huang, W. Malvidin Induces Hepatic Stellate Cell Apoptosis via the Endoplasmic Reticulum Stress Pathway and Mitochondrial Pathway. Food Science and Nutrition 2020, 8, doi:10.1002/fsn3.1810.

66. Saulite, L.; Jekabsons, K.; Klavins, M.; Muceniece, R.; Riekstina, U. Effects of Malvidin, Cyanidin and Delphinidin on Human Adipose Mesenchymal Stem Cell Differentiation into Adipocytes, Chondrocytes and Osteocytes. Phytomedicine 2019, 53, doi:10.1016/j.phymed.2018.09.029.

67. Huang, W.Y.; Liu, Y.M.; Wang, J.; Wang, X.N.; Li, C.Y. Anti-Inflammatory Effect of the Blueberry Anthocyanins Malvidin-3-Glucoside and Malvidin-3-Galactoside in Endothelial Cells. Molecules 2014, 19, doi:10.3390/molecules190812827.

68. Baba, A.B.; Nivetha, R.; Chattopadhyay, I.; Nagini, S. Blueberry and Malvidin Inhibit Cell Cycle Progression and Induce Mitochondrial-Mediated Apoptosis by Abrogating the JAK/STAT-3 Signalling Pathway. Food and Chemical Toxicology 2017, 109, doi:10.1016/j.fct.2017.09.054.

69. Mazewski, C.; Liang, K.; Gonzalez de Mejia, E. Comparison of the Effect of Chemical Composition of Anthocyanin-Rich Plant Extracts on Colon Cancer Cell Proliferation and Their Potential Mechanism of Action Using in Vitro, in Silico, and Biochemical Assays. Food Chemistry 2018, 242, doi:10.1016/j.foodchem.2017.09.086.

70. Castañeda-Ovando, A.; Pacheco-Hernández, M. de L.; Páez-Hernández, M.E.; Rodríguez, J.A.; Galán-Vidal, C.A. Chemical Studies of Anthocyanins: A Review. Food Chemistry 2009, 113.

71. Khoo, H.E.; Azlan, A.; Tang, S.T.; Lim, S.M. Anthocyanidins and Anthocyanins: Colored Pigments as Food, Pharmaceutical Ingredients, and the Potential Health Benefits. Food and Nutrition Research 2017, 61.

72. Burchmore, R.J.S.; Barrett, M.P. Life in Vacuoles – Nutrient Acquisition by Leishmania Amastigotes. International Journal for Parasitology 2001, 31, 1311–1320, doi:10.1016/S0020-7519(01)00259-4.

73. Moragas-Tellis, C.J.; Almeida-Souza, F.; do Socorro dos Santos Chagas, M.; de Souza, P.V.R.; Silva-Silva, J.V.; Ramos, Y.J.; de Lima Moreira, D.; da Silva Calabrese, K.; Behrens, M.D. The Influence of Anthocyanidin Profile on Antileishmanial Activity of Arrabidaea Chica Morphotypes. Molecules 2020, 25, doi:10.3390/molecules25153547.

74. Silva-Silva, J.V.; Moragas-Tellis, C.J.; Chagas, M.S.S.; Souza, P.V.R.; Moreira, D.L.; de Souza, C.S.F.; Teixeira, K.F.; Cenci, A.R.; de Oliveira, A.S.; Almeida-Souza, F.; et al. Carajurin: A Anthocyanidin from Arrabidaea Chica as a Potential Biological Marker of Antileishmanial Activity. Biomedicine and Pharmacotherapy 2021, 141, doi:10.1016/j.biopha.2021.111910.

75. Silva-Silva, J.V.; Moragas-Tellis, C.J.; Chagas, M.S.S.; Souza, P.V.R.; Moreira, D.L.; Hardoim, D.J.; Taniwaki, N.N.; Costa, V.F.A.; Bertho, A.L.; Brondani, D.; et al. Carajurin Induces Apoptosis in Leishmania Amazonensis Promastigotes through Reactive Oxygen Species Production and Mitochondrial Dysfunction. Pharmaceuticals 2022, 15, doi:10.3390/ph15030331.

76. Brust, R.; Lukacs, A.; Haigney, A.; Addison, K.; Gil, A.; Towrie, M.; Clark, I.P.; Greetham, G.M.; Tonge, P.J.; Meech, S.R. Proteins in Action: Femtosecond to Millisecond Structural Dynamics of a Photoactive Flavoprotein. J Am Chem Soc 2013, 135, 16168–16174, doi:10.1021/ja407265p.

77. Hünenberger, P.H.; Mark, A.E.; van Gunsteren, W.F. Fluctuation and Cross-Correlation Analysis of Protein Motions Observed in Nanosecond Molecular Dynamics Simulations. Journal of Molecular Biology 1995, 252, 492–503, doi:10.1006/jmbi.1995.0514.

78. Charlier, C.; Khan, S.N.; Marquardsen, T.; Pelupessy, P.; Reiss, V.; Sakellariou, D.; Bodenhausen, G.; Engelke, F.; Ferrage, F. Nanosecond Time Scale Motions in Proteins Revealed by High-Resolution NMR Relaxometry. J Am Chem Soc 2013, 135, 18665–18672, doi:10.1021/ja409820g.

79. Siqueira-Neto, J.L.; Moon, S.; Jang, J.; Yang, G.; Lee, C.; Moon, H.K.; Chatelain, E.; Genovesio, A.; Cechetto, J.; Freitas-Junior, L.H. An Image-Based High-Content Screening Assay for Compounds Targeting Intracellular Leishmania Donovani Amastigotes in Human Macrophages. PLoS Neglected Tropical Diseases 2012, 6, doi:10.1371/journal.pntd.0001671.

80. Rugani, J.N.; Quaresma, P.F.; Gontijo, C.F.; Soares, R.P.; Monte-Neto, R.L. Intraspecies Susceptibility of Leishmania (Viannia) Braziliensis to Antileishmanial Drugs: Antimony Resistance in Human Isolates from Atypical Lesions. Biomedicine & Pharmacotherapy 2018, 108, 1170–1180, doi:10.1016/j.biopha.2018.09.149.

81. Andrade, J.M.; Gonçalves, L.O.; Liarte, D.B.; Lima, D.A.; Guimarães, F.G.; de Melo Resende, D.; Santi, A.M.M.; de Oliveira, L.M.; Velloso, J.P.L.; Delfino, R.G.; et al. Comparative Transcriptomic Analysis of Antimony Resistant and Susceptible Leishmania Infantum Lines. Parasites and Vectors 2020, 13, doi:10.1186/s13071-020-04486-4.

82. Maes, L.; Beyers, J.; Mondelaers, A.; van den Kerkhof, M.; Eberhardt, E.; Caljon, G.; Hendrickx, S. In Vitro “time-to-Kill” Assay to Assess the Cidal Activity Dynamics of Current Reference Drugs against Leishmania Donovani and Leishmania Infantum. Journal of Antimicrobial Chemotherapy 2017, 72, doi:10.1093/jac/dkw409.

83. Oliveira, D.M.; Costa, M.A.F.; Chavez-Fumagalli, M. a.; Valadares, D.G.; Duarte, M.C.; Costa, L.E.; Martins, V.T.; Gomes, R.F.; Melo, M.N.; Soto, M.; et al. Evaluation of Parasitological and Immunological Parameters of Leishmania Chagasi Infection in BALB/c Mice Using Different Doses and Routes of Inoculation of Parasites. Parasitology Research 2012, 110, 1277–1285, doi:10.1007/s00436-011-2628-5.

84. Garg, R.; Dube, A. Animal Models for Vaccine Studies for Visceral Leishmaniasis. Indian Journal of Medical Research 2006, 123, 439–454.

85. Mears, E.R.; Modabber, F.; Don, R.; Johnson, G.E. A Review: The Current In Vivo Models for the Discovery and Utility of New Anti-Leishmanial Drugs Targeting Cutaneous Leishmaniasis. PLOS Neglected Tropical Diseases 2015, 9, e0003889, doi:10.1371/journal.pntd.0003889.

86. Pilon, A.C.; Valli, M.; Dametto, A.C.; Pinto, M.E.F.; Freire, R.T.; Castro-Gamboa, I.; Andricopulo, A.D.; Bolzani, V.S. NuBBEDB: An Updated Database to Uncover Chemical and Biological Information from Brazilian Biodiversity. Scientific Reports 2017, 7, 7215, doi:10.1038/s41598-017-07451-x.

87. Kim, S.; Chen, J.; Cheng, T.; Gindulyte, A.; He, J.; He, S.; Li, Q.; Shoemaker, B.A.; Thiessen, P.A.; Yu, B.; et al. PubChem 2019 Update: Improved Access to Chemical Data. Nucleic Acids Research 2019, 47, D1102–D1109, doi:10.1093/nar/gky1033.

88. Sander, T.; Freyss, J.; von Korff, M.; Rufener, C. DataWarrior: An Open-Source Program for Chemistry Aware Data Visualization and Analysis. Journal of Chemical Information and Modeling 2015, 55, doi:10.1021/ci500588j.

89. Daina, A.; Michielin, O.; Zoete, V. SwissADME: A Free Web Tool to Evaluate Pharmacokinetics, Drug-Likeness and Medicinal Chemistry Friendliness of Small Molecules. Scientific Reports 2017, 7, 42717, doi:10.1038/srep42717.

90. Bragina, M.E.; Daina, A.; Perez, M.A.S.; Michielin, O.; Zoete, V. The SwissSimilarity 2021 Web Tool: Novel Chemical Libraries and Additional Methods for an Enhanced Ligand-Based Virtual Screening Experience. International Journal of Molecular Sciences 2022, 23, doi:10.3390/ijms23020811.

91. Biasini, M.; Bienert, S.; Waterhouse, A.; Arnold, K.; Studer, G.; Schmidt, T.; Kiefer, F.; Cassarino, T.G.; Bertoni, M.; Bordoli, L.; et al. SWISS-MODEL: Modelling Protein Tertiary and Quaternary Structure Using Evolutionary Information. Nucleic Acids Research 2014, 42, 252–258, doi:10.1093/nar/gku340.

92. Dallakyan, S.; Olson, A.J. Small-Molecule Library Screening by Docking with PyRx. Methods in Molecular Biology 2015, 1263, doi:10.1007/978-1-4939-2269-7_19.

93. Trott, O.; Olson, A.J. AutoDock Vina: Improving the Speed and Accuracy of Docking with a New Scoring Function, Efficient Optimization, and Multithreading. Journal of Computational Chemistry 2009, 31, NA-NA, doi:10.1002/jcc.21334.

94. Hanwell, M.D.; Curtis, D.E.; Lonie, D.C.; Vandermeersch, T.; Zurek, E.; Hutchison, G.R. Avogadro: An Advanced Semantic Chemical Editor, Visualization, and Analysis Platform. J Cheminform 2012, 4, 1–17.

95. da Silva, A.W.S.; Vranken, W.F. ACPYPE-Antechamber Python Parser Interface. BMC Res Notes 2012, 5, 1–8.

96. Dolinsky, T.J.; Nielsen, J.E.; McCammon, J.A.; Baker, N.A. PDB2PQR: An Automated Pipeline for the Setup of Poisson–Boltzmann Electrostatics Calculations. Nucleic Acids Res 2004, 32, W665–W667.

97. van der Spoel, D.; Lindahl, E.; Hess, B.; Groenhof, G.; Mark, A.E.; Berendsen, H.J.C. GROMACS: Fast, Flexible, and Free. J Comput Chem 2005, 26, 1701–1718.

98. Case, D.A.; Belfon, K.; Ben-Shalom, I.; Brozell, S.R.; Cerutti, D.; Cheatham, T.; Cruzeiro, V.W.D.; Darden, T.; Duke, R.E.; Giambasu, G. Amber 2020. 2020.

99. Bussi, G.; Donadio, D.; Parrinello, M. Canonical Sampling through Velocity Rescaling. J Chem Phys 2007, 126, 014101.

100. Santos, K.B.; Guedes, I.A.; Karl, A.L.M.; Dardenne, L.E. Highly Flexible Ligand Docking: Benchmarking of the DockThor Program on the LEADS-PEP Protein–Peptide Data Set. J Chem Inf Model 2020, 60, 667–683.

101. Miller III, B.R.; McGee Jr, T.D.; Swails, J.M.; Homeyer, N.; Gohlke, H.; Roitberg, A.E. MMPBSA. Py: An Efficient Program for End-State Free Energy Calculations. J Chem Theory Comput 2012, 8, 3314–3321.

102. Valdés-Tresanco, M.S.; Valdés-Tresanco, M.E.; Valiente, P.A.; Moreno, E. Gmx_MMPBSA: A New Tool to Perform End-State Free Energy Calculations with GROMACS. Journal of Chemical Theory and Computation 2021, 17, 6281–6291.

103. Warnes, G.R.; Bolker, B.; Bonebakker, L.; Gentleman, R.; Liaw, W.H.A.; Lumley, T.; Maechler, M.; Magnusson, A.; Moeller, S.; Schwartz, M.; et al. Package “Gplots”: Various R Programming Tools for Plotting Data. R package version 2.17.0. 2016.

